# Mechanical regulation of Titin N2B-us conformation and its binding to FHL2

**DOI:** 10.1101/2022.09.05.506602

**Authors:** Yuze Sun, Wenmao Huang, Shimin Le, Jie Yan

## Abstract

The 572 amino acids unique sequence on titin N2B element (N2B-us) is known to regulate the passive elasticity of muscle as an elastic spring. It also serves as a hub for cardiac hypertrophic signaling by interacting with multiple proteins such as FHL1(Sheikh *et al*, 2008), FHL2(Lange *et al*, 2002), and Erk2(Perkin *et al*, 2015). N2B-us is thought to be an intrinsically disordered region. In addition, N2B-us bears force; therefore, the functions of N2B-us are likely regulated by mechanical stretching. In the work, we investigated the conformation of N2B-us as well as its force-dependent interaction with FHL2 using a combination of AlphaFold2 predictions and single-molecule experimental validation. Surprisingly, a stable alpha/beta structural domain (~115 a.a.) was predicted and confirmed in N2B-us, which can be mechanically unfolded at forces greater than 5 pN. More than twenty FHL2 LIM domain binding sites were predicted to spread throughout N2B-us including the regions cryptic in the structural domain. Mechanosensitive binding of FHL2 to N2B-us is revealed in single-molecule manipulation experiments. Together, the results unveil several previously unknown aspects of the N2B-us conformations and its force-dependent interactions with FHL2, which provides new insights into the physiological functions of the force-bearing N2B-us region.

## Introduction

Titin is the largest protein (3.4-3.9 MDa) in the human body that spans from Z disk to the M band region of the sarcomere and has essential structural and signaling functions in contractile units(Maruyama *et al*, 1977). The gene of titin, TTN, is expressed in three main full-length isoforms in adults with N2B isoform and N2BA isoform in the heart, as well as the N2A isoform in skeletal muscles. Selective expression of different isoforms has a huge impact on cardiac functions(Fatkin & Huttner, 2017; Kellermayer *et al*, 2021). There is an elastic spring-like part of titin in the I-band of the sarcomere, which connects thick filament and Z-disk. It acts as a molecular spring that is responsible for the assembly and maintenance of the sarcomere ultrastructure. This part of titin is composed of tandem Immunoglobulin (Ig) domain repeats and three intrinsically disordered regions, which are the PEVK region, N2A-us region, and N2B-us region. The three intrinsically disordered regions play differential roles in regulating titin functions in sarcomere(Linke *et al*, 1999; Zhou *et al*, 2016; Zhou *et al*, 2021).

Despite the mechanism remaining unknown, the N2B element plays an important role in the pathogenesis of cardiac diseases. Targeted deletion of N2B element in mouse leads to diastolic dysfunction and cardiac atrophy(Radke *et al*, 2007), and N2B-KO mouse has been used widely as a research animal model for cardiac diseases(Lee *et al*, 2013; Lee *et al*, 2010; Nedrud *et al*, 2011).

N2B-us was commonly considered as intrinsically disordered and serving as an elastic spring(Helmes *et al*, 1999). Its flexibility is highly modulated through phosphorylation by ERK2, CaMKIIδ(Perkin *et al*., 2015), PKG(Krüger *et al*, 2009), and PKA(Yamasaki *et al*, 2002). Consistent with this role, removal of the N2B-us was predicted to significantly increase the passive tension of titin (Pang *et al*, 2018). As a flexible force-bearing element in titin, the conformation of N2B-us is likely regulated by forces in the piconewton (pN) range(Linke *et al*, 1998), a range inaccessible to previous efforts to probe the force-dependent conformations of N2B-us using atomic force microscopy (AFM)(Zhu *et al*, 2009).

Further, N2B-us was found to interact with a bulk of signaling proteins including Erk2(Raskin *et al*, 2012), FHL1(Sheikh *et al*., 2008), FHL2(Lange *et al*., 2002), and *α*/*β*-crystallin(Zhu *et al*., 2009) and serves as a role of signaling hub thus having a great effect on cardiomyocytes functions. Dysregulated interactions between these factors to N2B-us are linked to diseases such as cardiac hypertrophy(van der Pijl *et al*, 2021). Being a force-bearing element in titin, it is reasonable to hypothesize that the interactions between N2B-us and these signaling proteins are regulated by mechanical stretching of N2B-us during heart beating. The force dependence of the interactions between N2B-us and most of its binding partners remains largely unexplored.

Among these N2B-us interacting signaling proteins, FHL2 (four and a half LIM protein-2) is a critical protein abundantly expressed in the heart, which has profound impacts on heart function. Mutations of FHL2 are associated with human cardiac hypertrophy(Friedrich *et al*, 2014). FHL2-deficient mice show no abnormality in heart development but have a serious lack of ability to recover from various types of cardiac insults(Hojayev *et al*, 2012; Kong *et al*, 2001). FHL2 binds to N2B-us, acting as a scaffold to recruit metabolic enzymes to titin(Lange *et al*., 2002) and negatively regulate ERK2(Purcell *et al*, 2004). In addition, FHL2 plays an important role in regulating the expression of several genes in cardiac tissue(Chan *et al*, 2000). However, little is known about the FHL2 binding sites in N2B-us, the conformation of the bound complex, and most importantly, how the binding is regulated by physiologically relevant forces.

To address these questions, in this study, making use of a combination of AlphaFold2 structural prediction and single-molecule manipulation experimental validation, we identified a thermodynamically stable structured domain in N2B-us with a high folding energy of ~ −8 kcal/mol, which unfolds at forces slightly above 5 pN. Using the same approaches, twenty-four LIM domain binding sites of FHL2 were predicted by AlphaFold2, which spread throughout N2B-us, including eight in the identified structural domain. Extensive high-affinity, multivalent binding of FHL2 to N2B-us was observed in single-molecule experiments, which is highly regulated by mechanical stretching. The binding of FHL2 to N2B-us leads to multiple force-dependent conformations, including a primary extended conformation facilitated by mechanical stretching.

## Methods

### AlphaFold2 prediction and PDB file analysis

Protein sequences of Titin N2B-us and FHL2 were obtained from Uniport (Q8WZ42 and Q14192), the sequence for full length N2B-us is DMTDTPCKAKSTPEAPEDFPQTPLKGPAVEALDSEQEIATFVKDTILKAALITEENQQLSYEHIA KANELSSQLPLGAQELQSILEQDKLTPESTREFLCINGSIHFQPLKEPSPNLQLQIVQSQKTFSKEGILMPEEPETQAVLSDTEKIFPSAMSIEQINSLTV EPLKTLLAEP EGNYPQSSIEPPMHSYLTSVAEEVLSPKEKTVSDTNREQRVTL QKQEAQSALIL SQSLAEGHVESLQSPDVMISQVNYEPLVPSEHSCTEGGKILIESAN PLENAGQD SAVRIEEGKSLRFPLALEEKQVLLKEEHSDNVVMPPDQIIESKRE PVAIKKVQE VQGRDLLSKESLLSGIPEEQRLNLKIQICRALQAAVASEQPGLFSEW LRNIEKV EVEAVNITQEPRHIMCMYLVTSAKSVTEEVTIIIEDVDPQMANLK MELRDALC AIIYEEIDILTAEGPRIQQGAKTSLQEEMDSFSGSQKVEPITEPEVESK YLISTEE VSYFNVQSRVKYLDATPVTKGVASAVVSDEKQDESLKPSEEK EESSSESGTEEVATVKIQEAEGGLIKEDG. The sequence for N2B-us structural domain is IPEEQRLNL KIQICRALQAAVA SEQPGLFSEWLRNIEKVEV EAVNITQEPRHIMCMYLVTSAKSVT EEVTIIIEDVDPQM ANLKMELRD ALCAIIYEEIDILT AEGPRIQQGAKT. The sequence for FHL2 is MTERFDCHHCNESLFGK KYILREESP YCVVCFETLFANTCEECGKPI GCDCKDLSYKDRHWHEACFHCSQCRNSLVDKPFAAKEDQLLCTDCYSNEYSSKCQECKKTIMPGTRKMEYKGSWHETCFICHRCQQPIGTKSFIPKDNQNFCVPCYEKQHAMQCVQCKKPITTGGVTYREQPWHKECFVCTACRKQLSGQRFTARDDFAYCLNCFCDLYAKKCAGCTNPISGLGGTKYISFEERQWHNDCFNCKKCS LSLVGRGFLTERDDILCP DCGKDI

AlphaFold2 code were ran in google colab page (https://colab.research.google.com/github/sokrypton/ColabFold/blob/main/beta/AlphaFold2_advanced.ipynb), configurations setting are as follows: msa_method=mmseqs2, pair_msa=False, max_msa=512:1024, subsample_msa=True, num_relax=0, use_turbo=True, use_ptm=True, rank_by=pLDDT, num_models=5, num_samples=1, num_ensemble=1, max_recycles=3, tol=0, is_training=False, use_templates=False. The exported PDB file were visualized and analyzed in Pymol 2.4.1.

### Protein expression and purification

The pET151-6His-Avi-2I27-N2B-2I27-Spy plasmid was prepared as previously described(Pang *et al*., 2018). Truncation of N2B-us folding domain was obtained by PCR (Q5® High-Fidelity DNA Polymerase, M0491) and Q5® High-Fidelity DNA Polymerase (M0491S) reaction, experiments were performed by the manufacturer’s instructions. Deletion of N2B-us folding domain was accomplished by PCR and DPN1 (NEB, R0176S) reaction with subsequent KLD (NEB KLD Enzyme Mix, M0554S) reaction, procedures were as described in manufacturer’s protocol. The sequences of primers, which were from IDT RxnReady® Primer Pools, were listed in Table S1. All constructs were validated by full sequencing of the open reading frame (Axil Scientific) and expressed in E. coli BL21 strain and purified using His-tag Coolum. Protein purity was evaluated by SDS-polyacrylamide gel electrophoresis, and protein concentration was measured by absorbance at 280 nm. Recombinant GST-FHL2 protein was purchased from Abnova (H00002274-P01), which was expressed in Wheat Germ.

### Single-molecule manipulation

Single-molecule N2B-us stretching experiments were performed on homemade magnetic-tweezers. The channels for single-molecule manipulation experiments were prepared as follows: 20×32 mm^2^ coverslips were cleaned by ultrasonication in detergent, acetone and add water for 20min each, after being fully dried in the oven, the coverslips underwent 15min oxygen plasma treatment. Afterward, the coverslips were salinized by incubation in 1% (3-Aminopropyl) triethoxysilane (APTES) (*Sigma*) in methanol for 20 min and rinsed thoroughly with methanol and cured for 20 min in the oven for drying. To prevent disturbance when changing buffer in the channel, we add a microwell array to the coverslips as described in(Le *et al*, 2015), then covered by 18*18 mm^2^ clean coverslip (*Deckglaser*) with two parallel stripes. The laminar flow channel was further treated with 1% glutaraldehyde for 1h. After fully rinsed by PBS (Phosphate Buffered Saline, Research instruments, SH30256.01), 3 μm diameter amine-coated polystyrene beads were added to the channel and incubate for 20min. Spy catcher expressed in E coli BL21 and purified by His tag column. and incubate in a concentration of 200ug/ml overnight. Afterward, 1% BSA in PBS buffer was added to the channel and incubate overnight for blocking.

Spy and avi tagged N2B-us protein were diluted to 0.002 mg/ml and were introduced in the channel and incubated for 30 min before the introduction of 2.8um diameter streptavidin-coated paramagnetic beads (M270 streptavidin, Dynabeads), incubate for 20 min for forming tethers. The buffer for the single-molecule manipulation experiment is PBS with 1% BSA and 10mM DTT. During experiments, the force on each bead was determined based on their thermal fluctuations under force, with a relatively small error (~10%).

Raw extension data were recorded at a sampling rate of 200Hz, data visualization and analysis were performed in OriginPro 2019b. To improve the clarity of data representation, some figures were smoothed using the fast Fourier transform (FFT)-smooth function in OriginPro 2019b. The force-extension curves were obtained by staying in each force for 20 s and taking the average extension, data processing was performed by in-house written Python 3.8 code and ran in spyder (anaconda 3). The illustration was performed in GraphPad Prism 7.

## RESULTS

### N2B-us contains a stable structural domain

To probe possible structural domains in N2B-us, we input the 572 a.a. N2B-us sequence to AlphaFold2(Jumper *et al*, 2021). Surprisingly, a structural domain of about 115 a.a. was predicted, which contains three beta strands spanned between two alpha-helical regions (Figure S1 & Figure 1A). To verify the existence and localization of such a structural domain in N2B-us, we prepared a protein construct where the sequence of the N2B-us region is inserted between four well-characterized titin Ig 27th domains (I27, two repeats at each end) acting as a molecular spacer(Yuan *et al*, 2017). The N- and C- termini of the construct contain a biotinylated AviTag and a SpyTag(Zakeri *et al*, 2012), respectively, for specific tethering to a coverslip surface and a superparamagnetic microbead (M270, Invitrogen) (Figure 1B). Time-varying or constant external forces were applied to tethered protein constructs via the end-attached microbead using an in-house built magnetic-tweezers setup(Chen *et al*, 2011; Zhao *et al*, 2017). The height of the microbead was recorded in real-time at a nanometre resolution. Due to force balance, the tension in the molecule is the same as the applied force. In addition, a stepwise bead height change at a constant or a time-varying force is the same as the extension change of the molecule(Zhao *et al*., 2017). The force was calibrated based on extrapolation with a standard curve, which is associated with 10% relative error as described in our previous publications(Chen *et al*., 2011; Zhao *et al*., 2017). The N2B-us region contains six cysteine residues. 10 mM DTT was included in the buffer solution to prevent formation of disulfide bonds (Grützner *et al*, 2009).

**Figure 1.**
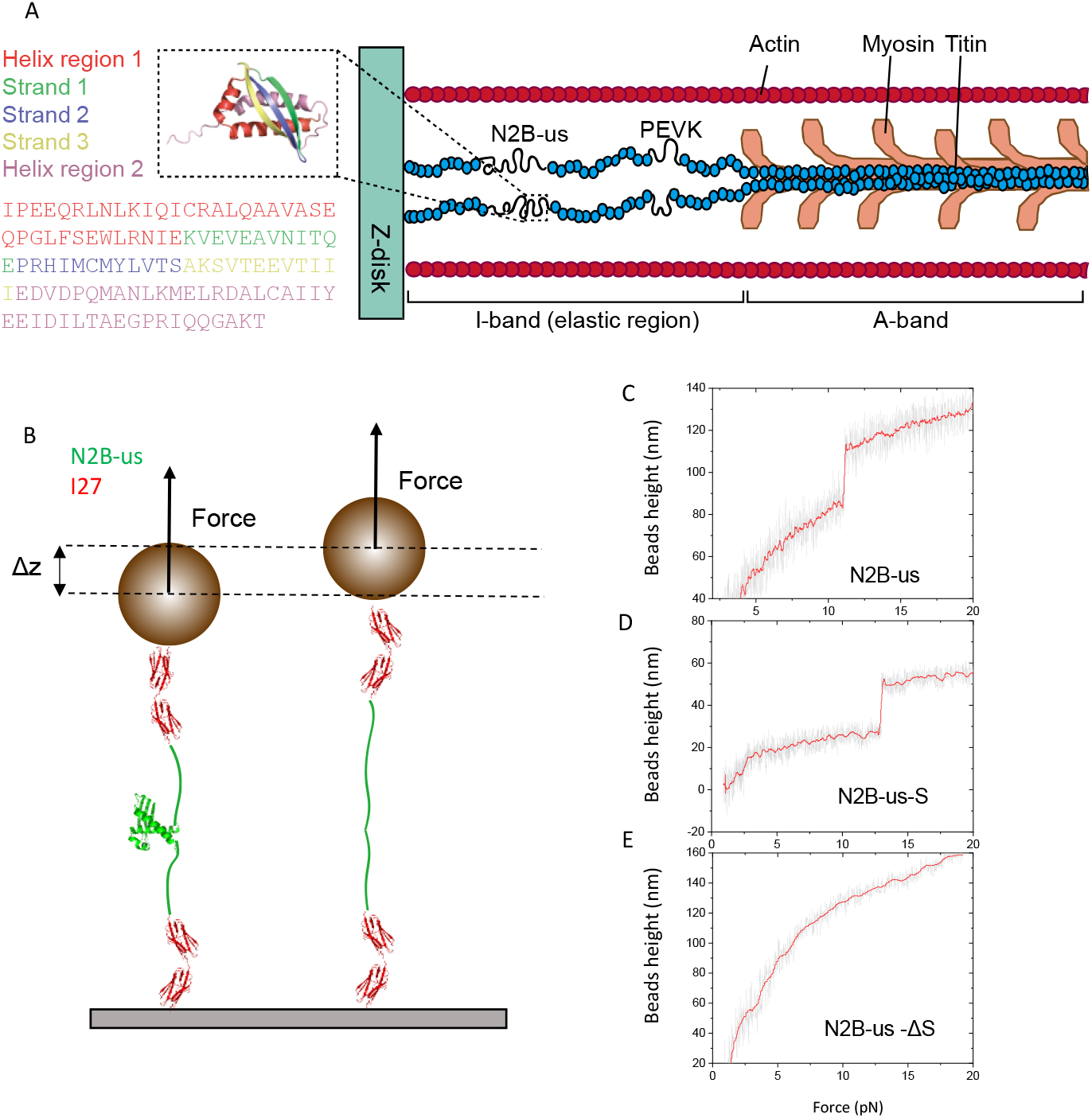
N2B-us structural prediction by Alphafold2 and single-molecule validation. **(A)** Schematic illustration of N2B-us position in sarcomere (right) and N2B-us structural domain (115a.a.) predicted by AlphaFold2 (left). The structure of the 115-amino-acid long domain contains three *β*-strands sandwiched between two *α*-helical regions, which are indicated by different colors. **(B)** Single-molecule detection of the conformations of N2B-us, the predicted structural domain (N2B-us-S), and N2B-us with the deletion of the structural domain (N2B-us-ΔS). The Illustration shows the protein construct for N2B-us tethered between a superparamagnetic microbead and a coverslip surface for an example. **(C-E)**. Representative time traces of the bead height during force-increase scans at a force loading rate of 1pN/s. The unfolding of the structured domain resulted in the stepwise height increases observed for the N2B-us and N2B-us-S constructs but not in N2B-us - ΔS.

Figure 1C shows a typical force-height curve of full-length N2B-us construct during a force-increase scan with a force-loading rate of 1 pN/s from 1pN. A stepwise bead height increase was observed at a force around 10 pN, indicating a stable compact element in N2B-us. To see whether this compact element corresponds to the AlphaFold2-predicted structural domain in N2B-us, a construct with only the sequence of 115a.a. structural domain (N2B-us-S) predicted by AlphaFold2 was expressed and purified. A similar stepwise bead height increase was observed when the experiment was repeated on N2B-us-S (Figure 1D). The force-dependent step size distribution is consistent with the force-dependent extension difference between the unfolded polypeptide chain and the folded structure (Figure 2 C, Text S2). Further, when the experiment was repeated on a construct containing the N2B-us sequence with the predicted structural domain deleted (N2B-us-ΔS), the stepwise bead height increase disappeared (Figure 1E). These results confirm the existence of the predicted structural domain in N2B-us, which could withstand a few pN forces.

**Figure 2.**
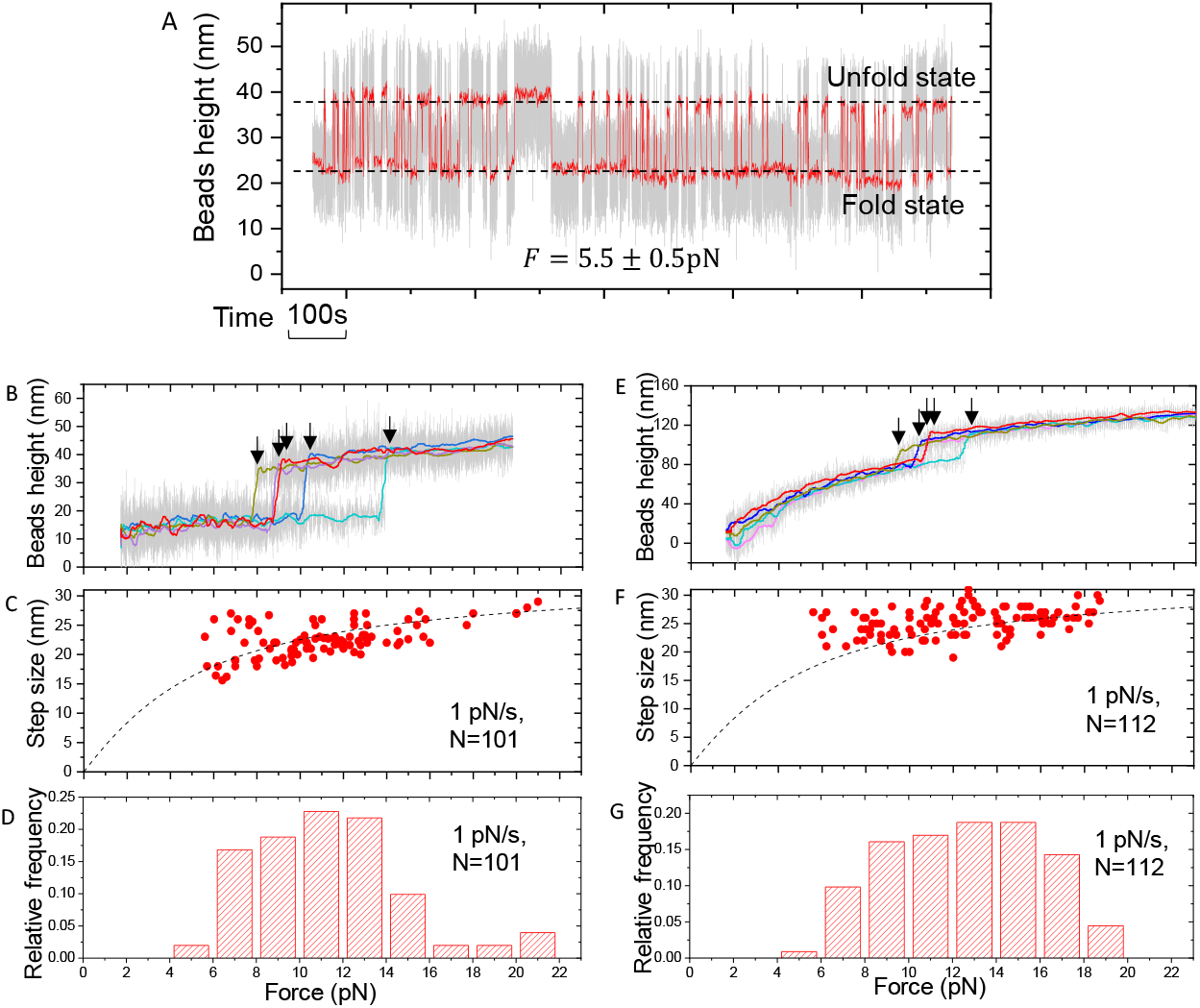
Dynamic transitions between the folded and unfolded states of N2B-us-S. **A).** A representative time trace of the height of a bead attached to a single protein construct containing an N2B-us-S at a constant force of 5.5 ± 0. pN. The stepwise changes of the bead height indicate spontaneous unfolding and refolding of the N2B-us-S, from which the overall folded-to-unfolded ratio can be obtained as the ratio of the fractions of time of the two states. **B).** Five representative force–height curves of N2B-us-S tether during force-increase scans at a loading rate of 1 pN/s. The coloured curves are obtained by 100-point FFT (fast Fourier transformation) smooth of the raw data (gray). The arrows indicate the unfolding events of N2B-us structural domain. **C-D).** The resulting force-dependent unfolding step sizes and the normalized force histogram obtained over 101 unfolding events from 5 independent tethers.**E-G)** unfolding of N2B-us structural domain under loading force in the full-length N2B-us construct. The unfolding force and step size are similar to that of N2B-us-S construct.

The structural domain underwent reversible unfolding and refolding fluctuations at a fixed force of *F* = 5.5 ± 0.5 pN (Figure 2A). The time fractions of the unfolded and the folded states are proportional to the probabilities of the corresponding states. The average ratio of the time fractions from data obtained from multiple independent tethers (Figure S4) with total time more than 3000s is determined to be 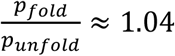, from which the zero-force folding energy was calculated to be Δ*G*_0_ = −13±1.91 K_B_T ≈ 8 kcal/mol by the equation 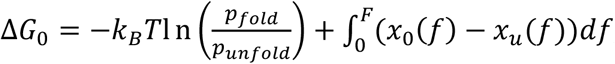, where *x*_0_(*f*) and *x*_*u*_(*f*) are the force-extension curves of the folded and unfolded states of the domain (Text S2). The error bar in the estimated Δ*G*_0_ was obtained by bootstrap analysis and error propagation calculation (Text S3). The absolute value of Δ*G*_0_ is comparable or greater than several known stable protein domains such as titin I27 Ig domain and *α* −actinin spectrin repeats(Yu *et al*, 2020). Together, these results confirmed that the predicted structural domain by AlphaFold2 is thermally stable, yet a mechanosensitive domain that can be unfolded by pN forces applied to N2B-us.

### FHL2 LIM domain binding sites spread throughout the N2B-us

N2B-us also plays a role in signal transduction by interacting with several signaling proteins including FHL2, a LIM domain protein involved in numerous cellular processes(Lange *et al*., 2002; Liang *et al*, 2018). An N2B-us region 79 a.a.−308 a.a. was reported to contain FHL2 binding sites(Lange *et al*., 2002), but the number of binding sites and their positions remain unknown. In addition, whether additional FHL2 binding sites exist outside the previously reported region is unclear. To address these questions, we applied AlphaFold2 to predict the structure of the complex of FHL2 and the subsegments within N2B-us and used a magnetic-tweezers-based single-molecule experiment to probe the interactions between N2B-us and FHL2.

AlphaFold2 predicts FHL2 binding to N2B-us with several conformations involving different binding sites in N2B-us (Figure 3A shows two representative structures of N2B-us – FHL2 complexes). A highly characteristic structure of the complex formed by an FHL2 LIM domain and a binding site on N2B-us is revealed: A single binding site (peptide of ~ 5 a.a.) on N2B-us antiparallelly associates with the first zinc finger in each FHL2 LIM domain via hydrogen bonding with the first *β*-strand in the zinc finger (Figure 3B). Dividing N2B-us into five subsegments each of ~100 a.a., FHL2 binding sites were predicted in each of the segments by AlphaFold2 (Figure S2A). Furthermore, AlphaFold2 predicted FHL2 LIM binding sites in each of the *β* -strands and the spanning alpha-helical regions in the structural domain (Figure S2B). In contrast, when the structural domain is folded, no binding was predicted (Figure S2C). These results suggest that FHL2 binding sites are spread throughout N2B-us, and some of them are cryptic in the folded structural domain. In total 24 FHL2 LIM domain binding sites were predicted, eight of which are buried in the folded structures (Figure 3C).

**Figure 3.**
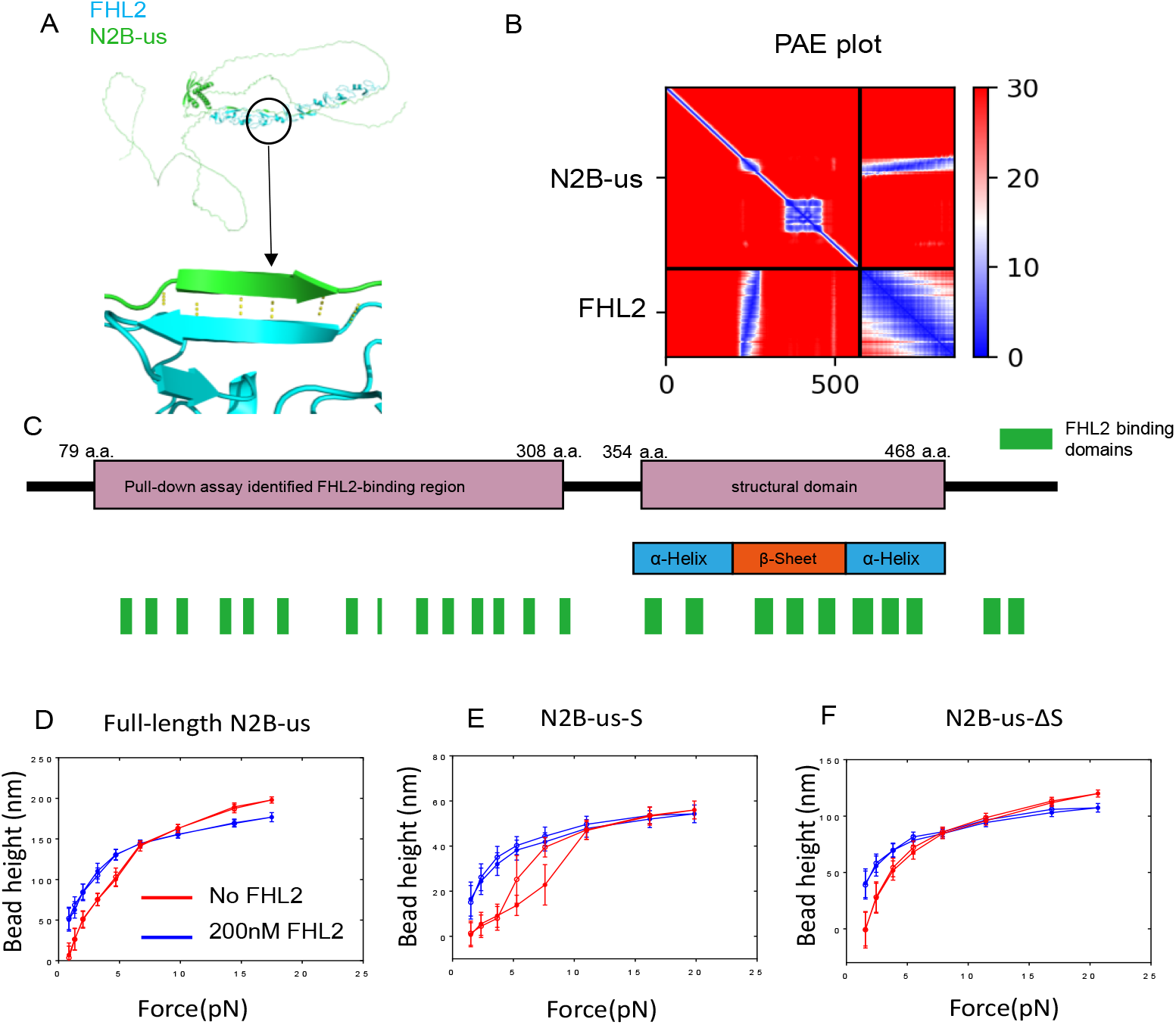
FHL2 binds to multiple sites on N2B-us. **(A)**. Structure of N2B-us in complex with FHL2. In one complex, the binding interface shows anti-parallel beta-strand conformation. **(B)**. predicted alignment error plot if the predicted structure. The binding interface shows low error (blue). **(C)**. FHL2-binding sites summary map for N2B-us. In total 23 FHL2-binding sites were predicted on N2B-us, 14 of which were in the previously reported segment identified by protein pull-down assay, 8 of which were cryptic in the structural domain. **(D)**. Representative force - height curves obtained from a full length N2B-us tether during force-increase (solid-circles) and force-decrease (open-circles) scans, before (red) and after (blue) introduction of 200 nM FHL2. The blue curves are higher than the red curves at forces below 8 pN. **(E)**. Representative force - height curves obtained from a N2B-us-S tether during force-increase (solid-circles) and force-decrease (open-circles) scans, before (red) and after (blue) introduction of 200 nM FHL2. Similarly higher blues curves than the red curves over pN force range was observed. **(F)**. Representative force -height curves obtained from a N2B-us-Δ*S* tether during force-increase (solid-circles) and force-decrease (open-circles) scans, before (red) and after (blue) introduction of 200 nM FHL2.

We tested the extensive distribution of the FHL2 LIM domain binding sites throughout N2B-us predicted by AlphaFold2 using magnetic-tweezers experiments. Figure 3D shows representative force-bead height curves recorded during a force-increase scan (solid circles) followed by a force-decrease scan (hollow circles) before (red) and after the (blue) introduction of 200 nM FHL2. The tether was held at each force for 20s, and beads height was obtained as the average value in the 20s. The results show a significant FHL2-dependent shift in the force-bead height curves, indicating FHL2 binding induced change of the N2B-us conformation. In this experiment, after the binding of FHL2, the height of the bead increased significantly at forces below 8 pN, while it decreased at forces above 8 pN. Similar, but different levels of a force-dependent shift in the force-bead height curves were observed from the same tethered molecule during different repeating cycles or from independent experiments under the same condition, suggesting the existence of different conformations of N2B-us/FHL2 complexes (Figure S3A).

We then tested the binding of FHL2 to the structural domain using the construct N2B-us-S. Before the introduction of FHL2, the representative force-bead height curve obtained during force-increase and force-decrease scans show large hysteresis, with the force-decrease curve higher than the force-increase curve. This is caused by the hysteric unfolding (occurred at higher forces during the force-increase scan) and refolding (occurred at lower forces during the force-decrease scan) of the structural domain. After introducing 200 nM FHL2 to the same tether, the force-bead height curves were significantly shifted, in a manner like that obtained from the full-length N2N-us construct (more representative data are provided in Figure S3B). In addition, the large hysteresis caused by domain unfolding and refolding disappeared.

We finally tested the binding of FHL2 to the region outside of the structural domain using the construct N2B-us-Δ S. Shifts in the force-bead height curves were also observed after the introduction of 200 nM FHL2, indicating FHL2 binding to this region of N2B-us which also caused a significant conformational change. More representative data are provided in Figure S3C.

Together, the results from AlphaFold2 prediction and the single-molecule binding assay show the existence of multiple FHL2 binding sites throughout N2B-us, which may organize N2B-us into different conformations indicated by different extensions at the same force. The binding-induced increased force-bead height under low force (~1pN) obtained from the N2B-us-ΔS construct suggests that FHL2 binding leads to a more extended conformation of the unstructured region. The loss of refolding of the structural domain observed from the N2B-us-S construct indicates a stably bound FHL2 that inhibits the domain refolding at low forces. In principle, as the prediction shown in Figure 3A (bottom), the four and a half domains on FHL2 may allow one FHL2 to engage distal FHL2 LIM domain binding sites on N2B-us resulting in a highly compacted conformation, which was confirmed in our experiment by holding the N2B-us-ΔS construct at a lower FHL2 concentration (10 nM) at a low force of 1 pN for 1000s (Figure S3D).

### Mechanical activation of FHL2 binding to the N2B-us structural domain

The results obtained from the N2B-us-S construct demonstrate the binding of FHL2 that subsequently suppresses the refolding of the structural domain at lower forces. We also showed that the folded N2B-us-S is thermally stable and requires >5 pN to unfold. While these results strongly suggest that mechanical unfolding of N2B-us-S may be necessary to expose the FHL2 LIM domain binding sites for high-affinity binding of FHL2, direct experimental evidence of this necessity is still lacking.

To address this question, we probed any possible binding of folded N2B-us structural domain to FHL2. In the experiments, FHL2 (200nM) was added to a folded N2B-us-S construct. Then, the time-increase force was applied at a loading rate of 1 pN/s from 1pN to 6pN. The upper limit of the force was chosen to avoid force-induced rapid unfolding of the N2B-us-S structure. If FHL2 could invade and gain access to the FHL2 LIM domain binding sites in the structural domain, one would expect FLH2-dependent unfolding of the domain over the low force range, which can be determined based on the change of the bead height. Figure 4A shows that the bead height remained at a level corresponding to the folded state of the N2B-us-S more than ten force cycles (two are shown in figure 4A and more can be found in figure S6). In contrast, when the upper bound of the scanned force range was increased to 22 pN, stable binding of FHL2 occurred after the first force-increase cycle during the time when the N2B-us-S domain was in the unfolded state (Figure 4B). The binding was indicated by loss of refolding of the domain during the force-decrease scan at a loading rate of −1 pN/s (the dashed red line). During the subsequence force-increase and force-decrease scans, the bead height further increased, indicating more FHL2 binding or additional change of the configuration of the bound complex.

**Figure 4.**
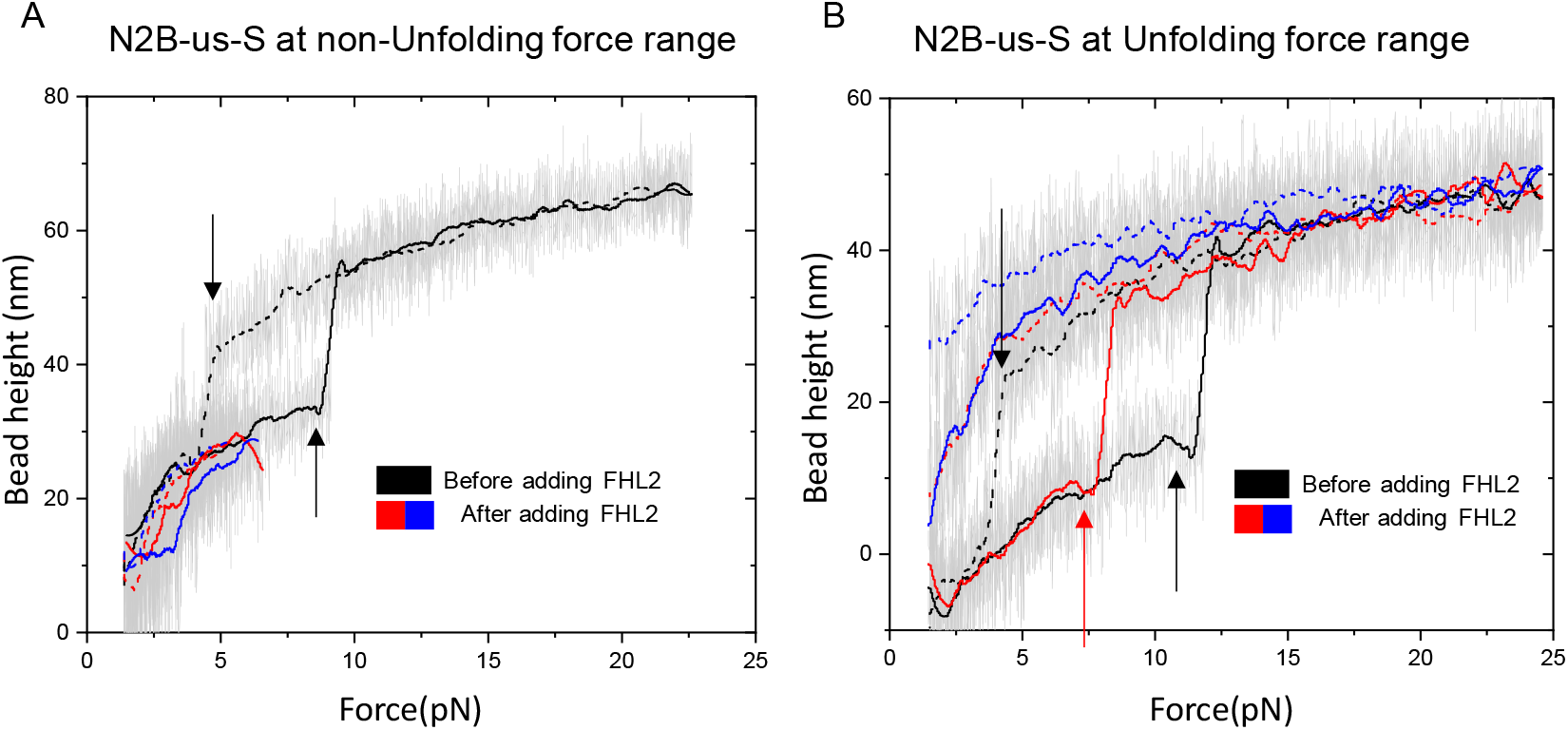
Force-activated FHL2 binding to the N2B-us structural domain. **(A)**. Representative force-bead height curves of N2B-us-S obtained during force-increase (solid lines) and decrease (dashed lines) scans, before (black lines) and after (colored lines) adding 200 nM FHL2. The lines represent 100-point FFT smoothing from the raw data (gray). Before adding FHL2, the force-dependent unfolding and refolding steps of N2B-us structural domain are indicated by red arrows. After adding FHL2, the scanned force range was capped at 6 pN to avoid rapid force-dependent unfolding during the scan. The data show that the N2B-us structural domain remained folded in 200 nM FHL2. **(B)**. Repeating the same experiments with increased upper limit of the scanned force range to 25 pN, after unfolding of the structural domain in 200 nM FHL2 in the first force-increase scan, the refolding signal was lost during the subsequent force-decrease scan. In all following force-increase and force-decrease cycles, the force-bead height curves correspond to the unfolded state of the structural domain.

Together, the results indicate that FHL2 cannot spontaneously invade into N2B-us-S in the folded state at low forces, and its high-affinity binding to FHL2 strictly requires mechanical activation.

## DISCUSSION

In summary, using an integrated approach that combines AlphaFold2 structural and binding predictions and single-molecule manipulation approach, our studies have revealed a mechanosensitive structural domain in the N2B-us region, and a plethora of FHL2 LIM domain binding sites in N2B-us that leads to mechanosensitive, high-affinity multivalent binding to FHL2. Figure 5 illustrates different configurations of the FHL2-N2B-us complex formed at different forces.

**Figure 5.**
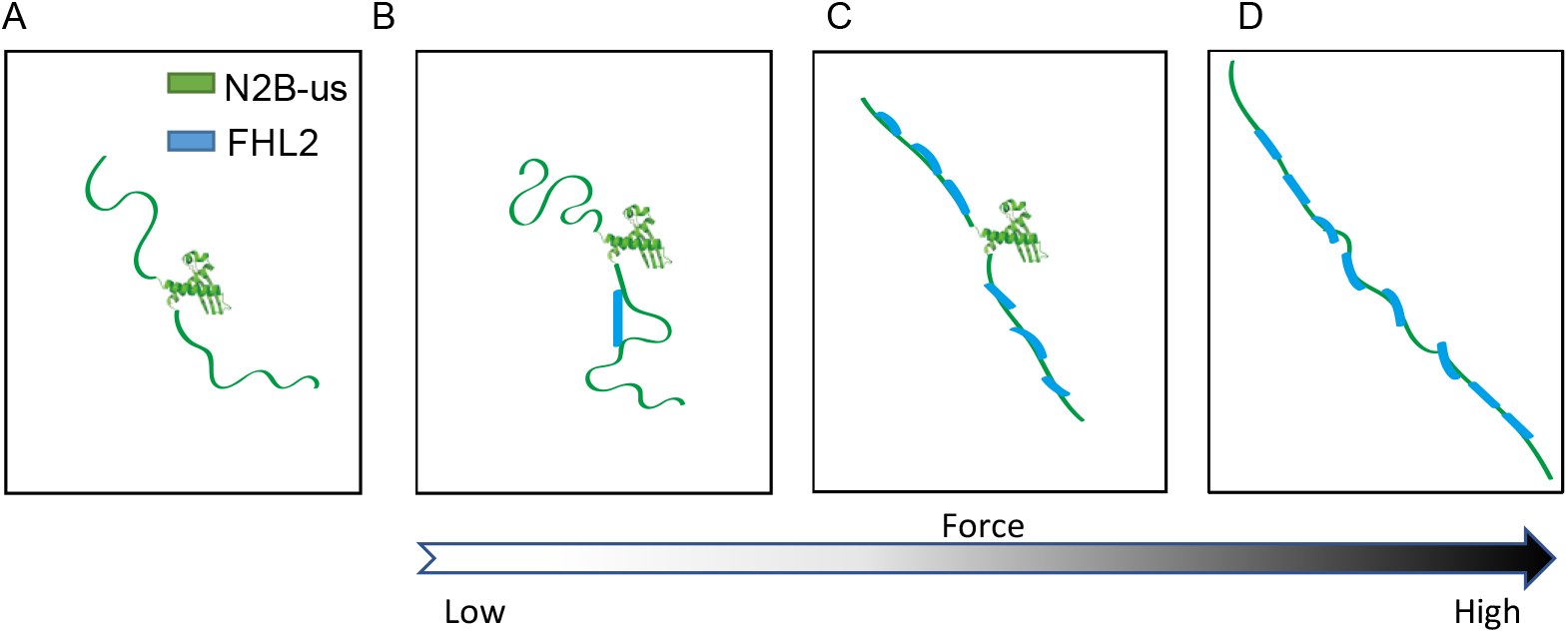
Schematic diagram of force-dependent configurations of the FHL2-N2B-us complexes. **(A)** unbound N2B-us. **(B)** FHL2-N2B-us complex formed only in low FHL2 concentration and at low forces where the randomly coiled unstructured region promotes binding of FHL2 to distal FHL2 LIM domain binding sites resulting in N2B-us looping. **(C)** FHL2-N2B-us complex at medium forces where the more extended unstructured region promotes binding of FHL2 to adjacent FHL2 LIM domain binding sites resulting in stiffening of N2B-us. **(D)** FHL2-N2B-us complex at high forces where the structural domain is unfolded, and the extended N2B-us promotes binding of FHL2 to adjacent FHL2 LIM domain binding sites resulting in stiffening of N2B-us. These configurations, once formed, can persist during cyclic heart beating.

N2B-us has long been thought to be an intrinsically disordered region. Surprisingly, AlphaFold2 predicts a well-folded structural domain (~115 a.a.) in N2B-us. It was successfully purified and confirmed to be a thermodynamically stable domain using a single-molecule manipulation approach. In our previous study, it was found that N2B-us exhibited a collapsed conformation at forces below 5 pN (Pang *et al*., 2018). The structural domain identified in this study provides an explanation for that observation. Therefore, our results show that the 574 a.a. of N2B-us is a mixture of a ~460 a.a. unstructured randomly coiled peptide region and a 115 a.a. structural domain.

N2B-us is a force-bearing element in titin, which is subject to periodic change of tension in the pN force range during heart beating(Linke *et al*., 1998; Pang *et al*., 2018). The extension of the ~460 a.a. unstructured region is sensitive to force change over several pN force range. Tensile forces of 6-9 pN are needed to stretch the region to half of its contour length based on the typical peptide polymer bending persistence length range of 0.5 – 0.8 nM (Chen *et al*, 2015; Winardhi *et al*, 2016) (Text S2). The ~ 115 a.a. structural domain is unfolded to a peptide polymer at forces above ~5 pN of constant forces. These results indicate that both the disordered region and the structural domain in N2B-us are sensitive to physiological levels of tension; therefore, the conformation of N2B-us is sensitively regulated by tension change during heart beating cycles.

Besides acting as an elastic spring to regulate the passive tension in titin, N2B-us also plays a role in signal transduction by interacting with several signaling proteins including FHL2. Previous pull-down assay suggested that N2B-us interacts with FHL2 via a region of 79a.a.-308a.a (Lange *et al*., 2002). However, the exact FHL2 binding sites on N2B-us were not identified. In this work, AlphaFold2 predicted twenty-three FHL2 LIM domain binding sites, each being a linear peptide of 3-5 a.a. Thirteen sites are clustered in a disordered region from 79a.a-308a.a., and eight are clustered and buried in the folded structure. In each cluster, the FHL2 LIM domain binding sites are linked by short peptides (5-15 a.a.). AlphaFold2 also predicted a highly characteristic structure of the complex formed by an FHL2 LIM domain and a binding site on N2B-us. A single binding site on N2B-us antiparallelly associates with the first zinc finger in each FHL2 LIM domain via hydrogen bonding with the first *β*-strand in the zinc finger, resulting in an extended conformation.

Consistent with these predicted binding sites, our single-molecule experiments revealed high-affinity binding of FHL2 to both the disordered region and the region containing the structural domain. The binding to the disordered region results in slightly increased bead height in pN force range. The shift in the force-bead height curves varied significantly in different experiments, which can be explained by different configurations of FHL2 binding to N2B-us. FHL2 has four and half LIM domains, which can engage different set of binding sites on N2B-us, resulting in many potential bound configurations associated with different force-dependent extensions. The binding to the exposed binding sites in unfolded structural domain kinetically inhibits refolding of the structural domain when forces were decreased to below 5 pN. The numerous FHL2 binding sites make the N2B-us a hot spot for FHL2 binding, which could scaffold downstream binding and potential activation of several important kinases such MAPK and PFK(van der Pijl *et al*., 2021).

The force-dependent conformations of N2B-us imply mechanosensitive interactions between N2B-us and FHL2. Importantly, force-dependent binding of FHL2 to N2B-us has been proposed to play a crucial role in regulating cardiac hypertrophy and atrophy(Radke *et al*, 2019). However, whether mechanosensitive interaction between N2B-us and FHL2 exists, or the underlying potential mechanism remain unclear. We showed that the binding sites buried in the structural domain required mechanical unfolding for exposure, which is activated for binding to FHL2 by forces greater than 5 pN. The large folding energy of ~ − 8 kcal/mol of the structural domain suggests that the binding affinity to FHL2 could be increased by five orders of magnitude upon mechanical unfolding of the domain. The FHL2 binding sites in the intrinsically disordered region are also mechanosensitive. The clustered FHL2 LIM domains are arranged as a linear array with an inter-domain distance of similar lengths (~3 nm), making it energetically favorable for FHL2 to bind several adjacent FHL2 LIM domain binding sites on N2B-us. Our single-molecule manipulation assay showed that the N2B-us/FHL2 complex took a more rigid, extended conformation compared to naked N2B-us; hence, the affinity of FHL2 binding to neighboring sites on N2B-us is predicted to be increased by tensile forces of a few pN (Efremov & Yan, 2018; Yan & Marko, 2003) (Text S1). Together, the results suggest a two-tier mechanosensitive binding between N2B-us and FHL2.

One proposed functional role of N2B-us is that it acts as a passive force buffer due to the highly flexible nature of randomly coiled peptide polymer. This picture needs revision based on the discovered stable structural domain within N2B-us and its numerous binding sites for FHL2. Below a threshold force ~5 pN, the structural domain absorbed ~1/5 of the residues in N2B-us, resulting in a tighter conformation than previously thought. When the domain unfolds, the released additional flexible peptide will reduce the tension. The extensive FHL2 binding to N2B-us can also impact the force-buffer role of N2B-us. The FHL2 binding induced rigid, extended conformation of N2B-us could effectively reduce the tension level across titin.

This study demonstrates a powerful integration of AlphaFold2 prediction and single-molecule experiments to reveal the structure-interaction relationship of proteins whose structures have not been solved experimentally. Single-molecule mechanical manipulation can provide useful information on the dynamic protein conformations and interactions. Much deeper insights can be obtained if the structural information of the proteins and the protein-protein complexes can be integrated into the interpretation of the results from the single-molecule manipulation studies. In this study, AlphaFold2 predicts a previously unknown structural domain in titin N2B-us, numerous FHL2 LIM domain sites in N2B-us, and the conformation of the complexes formed between N2B-us and FHL2, which form the structural basis to understand the observed force-dependent conformational change of N2B-us and the mechanosensitive interaction between N2B-us and FHL2 from our single-molecule mechanical manipulation experiments. While the predictions from AlphaFold2 require future structural biological studies to confirm, the predicted structural domain in N2B-us and the FHL2 binding regions are fully consistent with the results from our single-molecule manipulation experiments. We believe that the combination of AlphaFold2 and single-molecule manipulation will become as a powerful generic methodology to understand the relationship between the structures and dynamic changes of conformation and interactions of a broad range of proteins.

