## Supplementary material for "Mechanical regulation of Titin N2B-us conformation and its binding to FHL2": SI figures and text

Figure S1

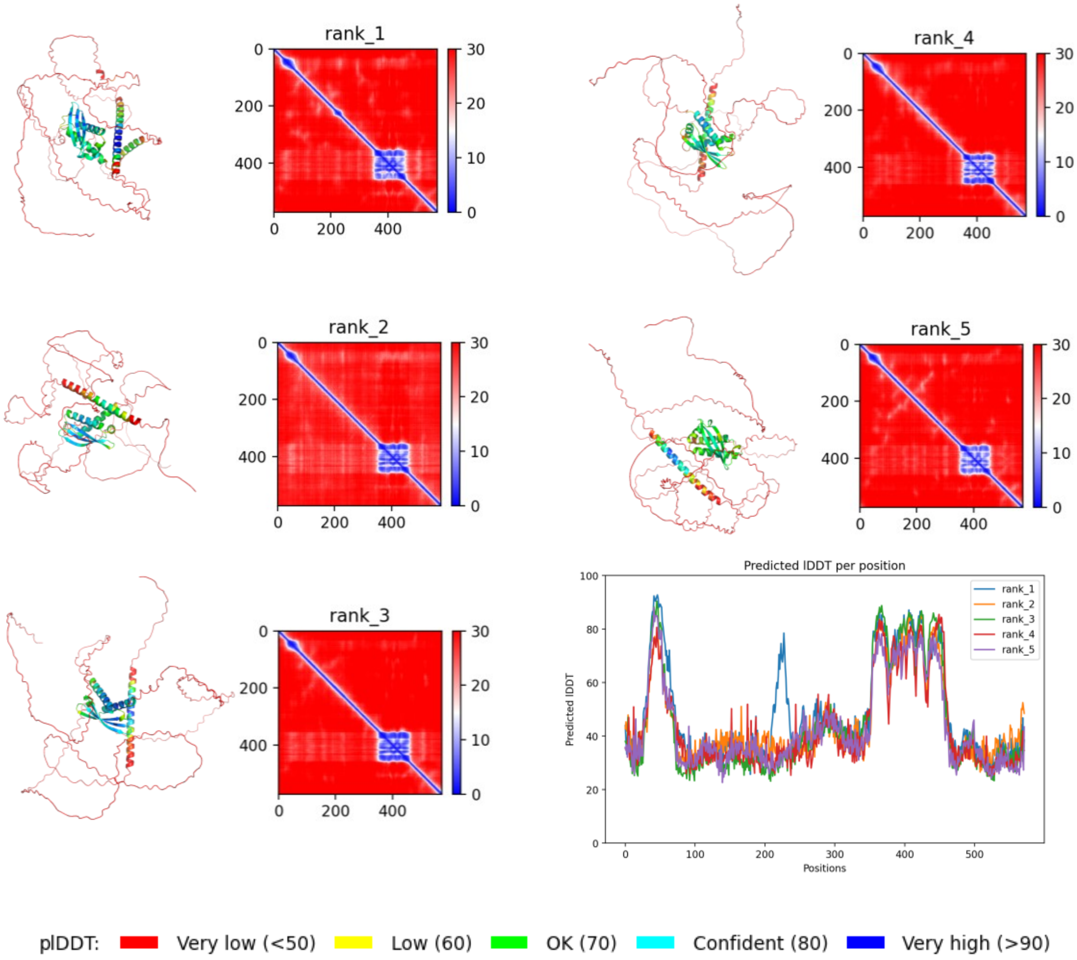


**Figure S1 AlphaFold2 predicts the structure of full length N2B-us.** A ~115a.a. structural domain is predicted in N2B-us in all the four given models, which are indicated by green color.

Figure S2

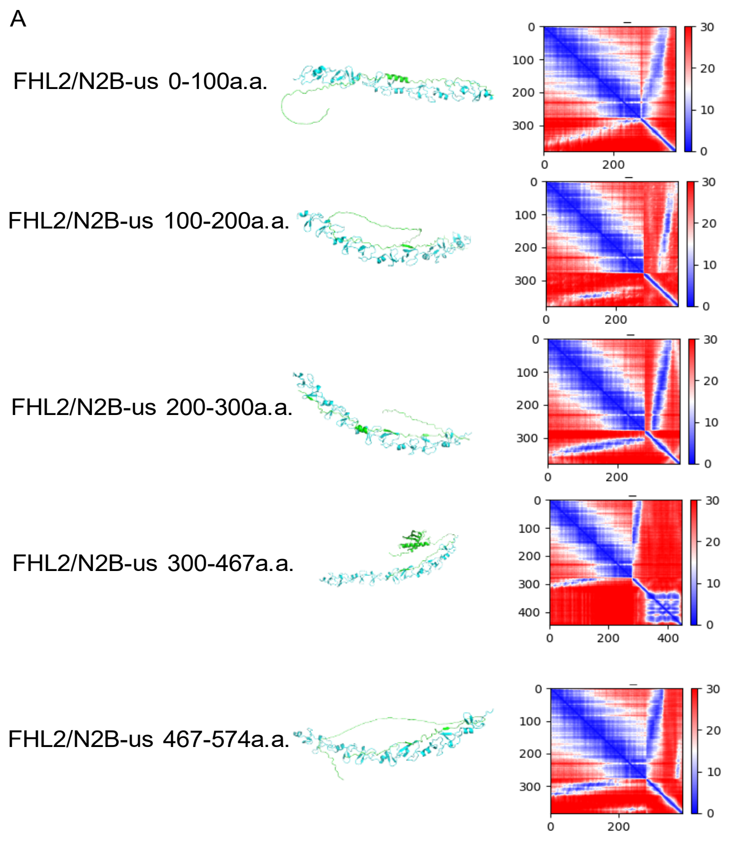


**Figure S2. Structure of FHL2/N2B-us fragment complex predicted by AlphaFold2 A).** Five subsegments of N2B-us in complex with FHL2, revealing FHL2 LIM domains binding sites in each of the segment. The structural domain is retaind in the segment of 300-467 a.a. **(B)** 115 a.a. long sequence of N2B-us structural domain and sequence of FHL2 were input to AlphaFold2. **(C).** AlphaFold2 predicts no binding between the structured domain and FHL2 (top).

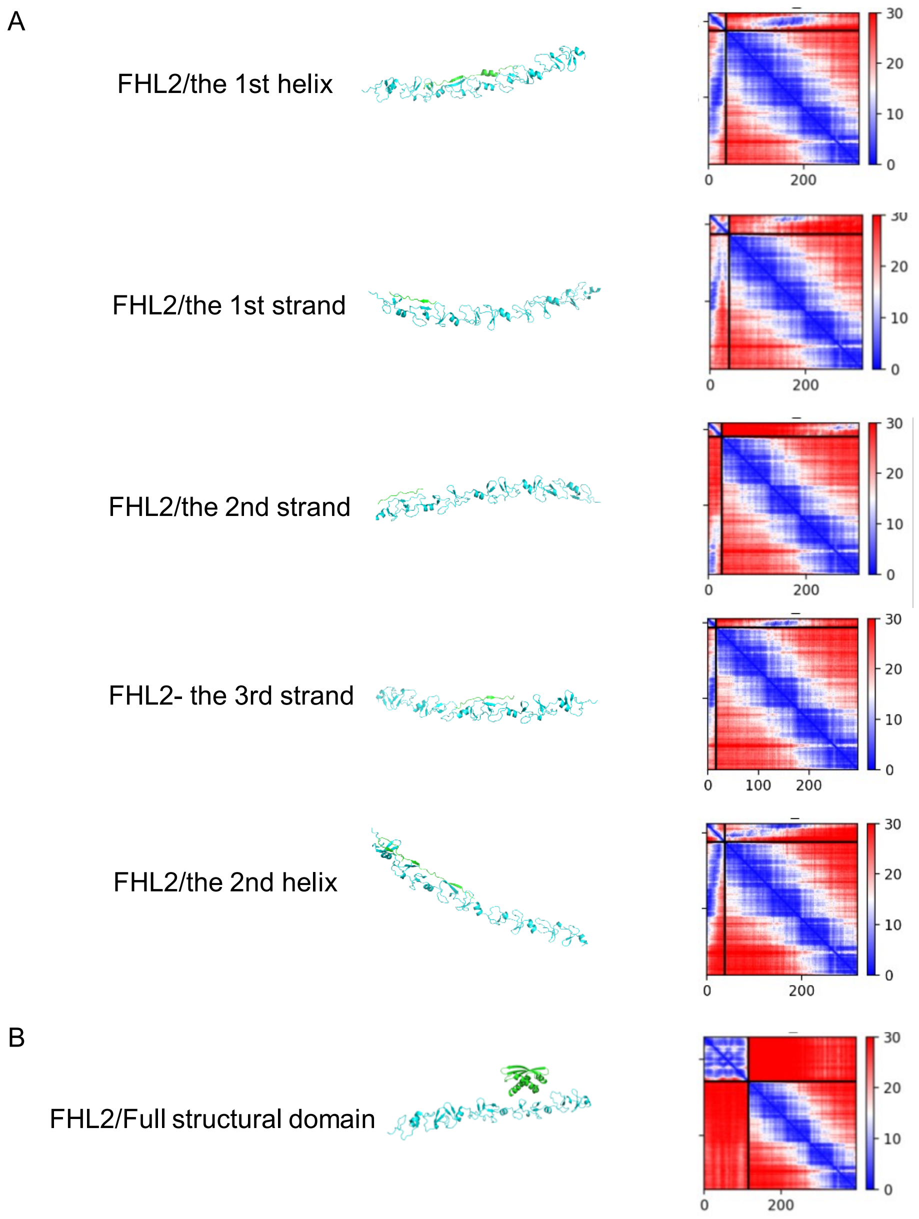


**Figure S3. Structure of FHL2/N2B-us structural domain fragments complex predicted by AlphaFold2 A).** The structure of each helical/strand region of N2B-us structural domain in complex with FHL2. On the right is the corresponding PAE plot. They all show confident binding sites. **(B)** The whole 115 a.a. long sequence of N2B-us structural domain in complex with FHL2. The structure and PAE plot show no binding.

Figure S3

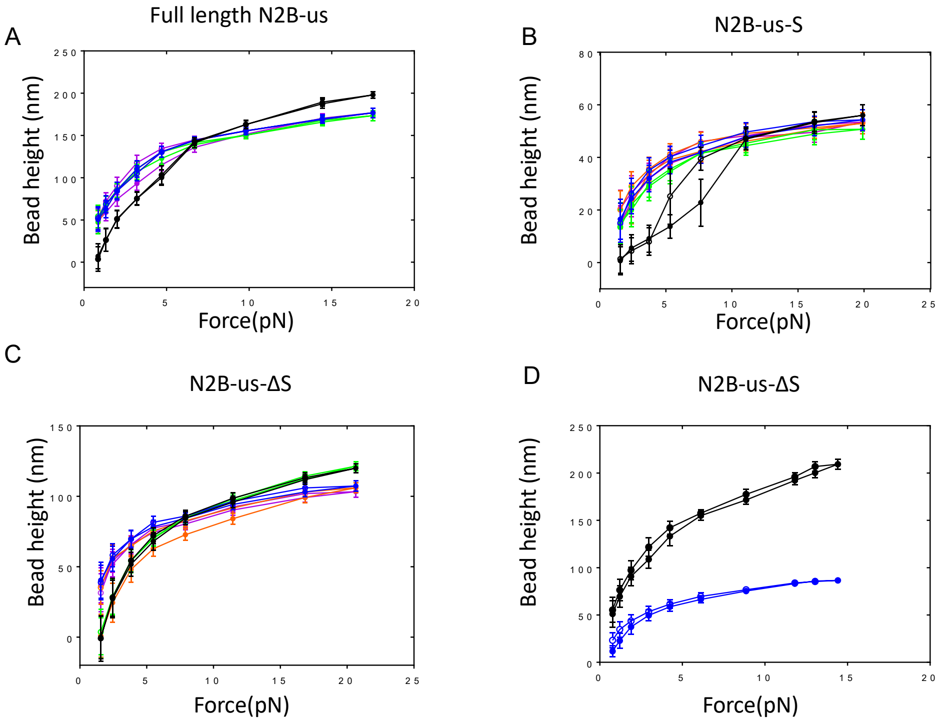


**Figure S3 Multiple cycles of force extension curves for N2B-us constructs binding with FHL2.** Black curve represents for the bead height before introducing FHL2, colored curves represent for bead height after introducing 200nM FHL2**. (A).** 1 cycle force-extension characterization of full length N2B-us without FHL2 and 4 cycles of force-extension of N2B-us in 200nM FHL2. **(B-C).** same characterization for N2B-us-S and N2B-us-ΔS constructs. **(D).** An extreme case of FHL2 induced looping of N2B-us. In a N2B-us-ΔS construct, after low concentration (10nM) FHL2 introduced and wait for more than 1000s at 1pN force, the height of bead dramatically decreased.

Figure S4

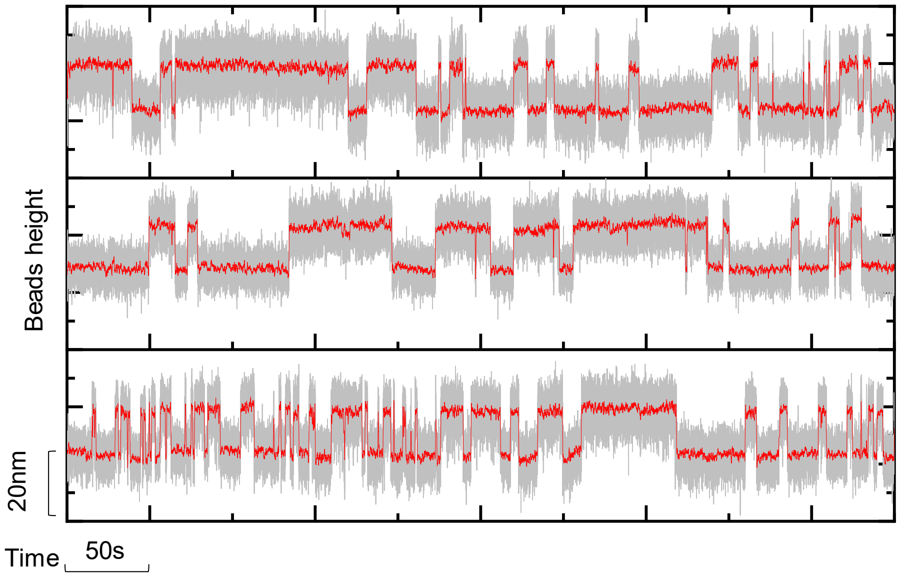


**Figure S4 Three tethers of N2B-us-S construct under 5.5pN undergo unfold and refold.** Red line is 100 FFT transformation smooth form the original data in grey. Beads heigh goes up and down with 15nm long step indicate the unfolding and refolding of the structural domain.

Figure S5

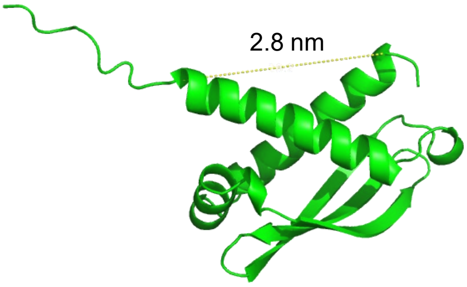


**Figure S5 Measurement of N- to C- terminal distance of folded N2B-us structural domain based on the prediction of AlphaFold2.** PDB file was obtained from AlphaFold2 prediction based on 115 a.a. long N2B-us structural domain sequence. The measurement is from 356E to 458E, the distance is ~2.8nm.

Figure S6

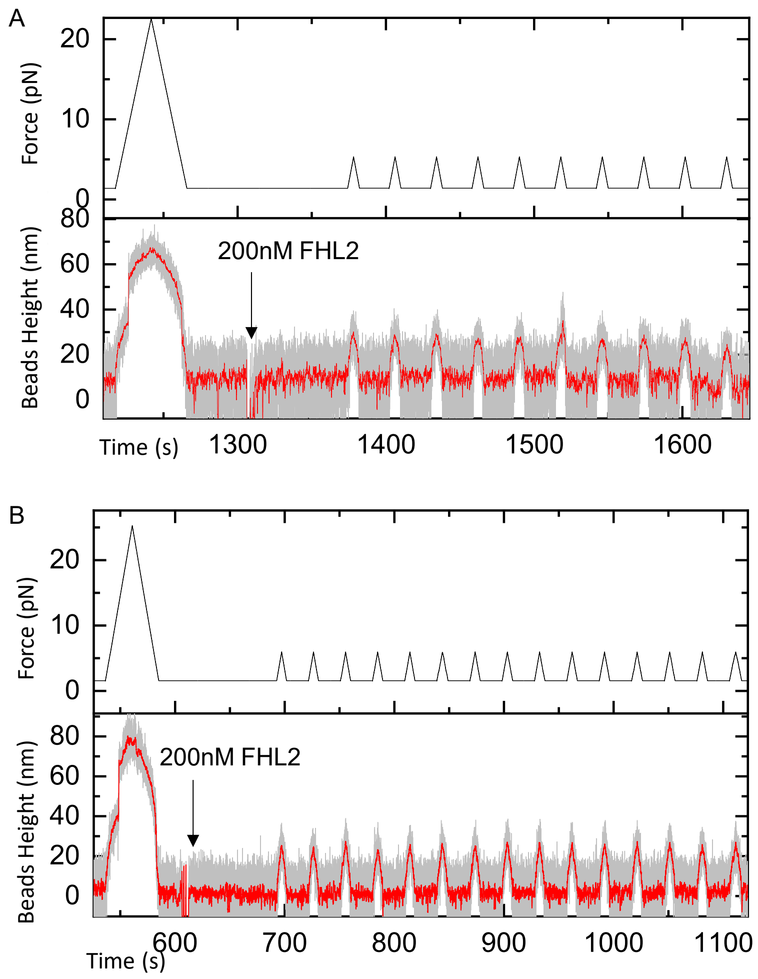


**Figure S6 loading force from 1-6pN for N2B-us-S in 200 nM FHL2. (A).** After 1 cycle of loading force from 1pN to 22pN and back to 1pN with loading rates 1pN/s and -1pN/s, respectively, 200 nM FHL2 was add to the buffer as indicated by the arrow. Subsequent loading forces from 1pN to 6pN didn't induce any binding of FHL2 to N2B-us. **(B).** Similar results in another bead.

### SI Text:

#### Text S1: The affinity of FHL2 binding to neighboring sites on N2B-us is predicted to be increased by theoretical prediction

The predicted FHL2 LIM domain binding sites are clustered into two regions, a region from 79 – 308 a.a. containing 13 sites and the other (354 – 468a.a.) corresponding to the folded structures, linked by short peptide (5-15 a.a.) of a few nm. Interestingly, the

LIM domains FHL2 is arranged as a linear array with an inter-domain distance of similar lengths (~3 nm). Therefore, it is energetically preferable for FHL2 to bind adjacent FHL2 LIM domain binding sites in N2B-us, resulting in a rigid, extended conformation of N2B-us/FHL2 complex. By maintaining a pre-extended conformation the unstructured region of N2B-us, forces of a few pN should significantly increase the binding affinity between FHL2 and N2B-us, with a force-dependent dissociation constant of $K_d(F) = K_d^0 e^{-\Delta\Phi(F)/k_B T}$, where $K_d^0$ is the dissociation constant at zero force, $k_B$ is the Boltzmann constant, $T$ is the temperature in the Kelvin unit, and $\Delta\Phi(F) = -\int_0^F \Delta x(f) df$ is the force-dependent conformational free energy difference between the unbound and bound states of N2B-us where $\Delta x(f)$ is the force-dependent extension difference between the two states[1]. It can be estimated when force increases from zero to 10 pN, $K_d(F)$ can decrease by more than 10 times, suggesting a force-facilitated binding of FHL2 to the unstructured, FHL2 binding region. The other FHL2 binding region, namely the structural domain, inhibits FHL2 binding unless it is unfolded at forces above 10 pN. The large folding energy of $\sim 10\ k_B T$ of the structural domain makes it a highly efficient mechanical switch[1], which is switched on for FHL2 binding at forces greater than 10 pN, associated with an increased binding affinity by five order of magnitude. Together, these results suggest that the N2B-us binding by FHL2 is mechanically modulated in a graded manner.

**Text S2 Theoretical force-dependent step-size of N2B-us structural domain unfolding/rupturing transitions**

A folded domain or complex can be considered as a rigid body. Hence, the force-extension curve of a folded domain or complex is determined by the rigid rotation fluctuation of a rigid body with a characteristic length b ~ 2.8 nm (Figure S4), estimated from the PDB file predicted by AlphaFold2, which is the distance between the two force-attaching points on the stable folded core (yellow dotted line in Fig. S4, excluding the N- terminal 2 a.a. and C-terminal 10 a.a. predicted unstable for the isolated domain by AlphaFold2). The corresponding force-extension curve can be described by the freely jointed chain polymer model with a single segment: $x^{FJC}(F) = b\left(coth\left(\frac{Fb}{K_BT}\right) - \frac{K_BT}{Fb}\right)$, where $K_BT = 4.1\, pN \cdot nm$ at room temperature.

The force-dependent step sizes during unfolding of the domain can be described assuming the unfolded state a randomly disordered polypeptide chain, using the worm-like chain (WLC)[2] polymer model with a reasonable bending persistence length of A ~ 0.8 nm [3]: $\frac{FA}{K_BT} = \frac{1}{4\left(1-\frac{x^{WLC}(F)}{L}\right)^2} - \frac{1}{4} + \frac{x^{WLC}(F)}{L}$ . Here $L = n * 0.38$ nm is the contour length of the unfolded state, where $n$ is the number of residues of the folded core (103 a.a.). The force-dependent unfolding/rupturing step size is the extension differences of the domain between the unfolded and folded states at the transition (unfolding/refolding) force, i.e.,$\Delta x(F) = x^{WLC}(F) - x^{FJC}(F)$. A catachrestic force of $\sim\frac{k_BT}{A}$ is needed to extend a randomly coiled peptide polymer to half of its contour length. Over the typical bending persistence length range, 0.5 nm – 0.8 nm[4,5] of peptide polymer, the half-contour length-extension tensile force is over a range of 5-8 pN.

**Text S3 Error estimation for folding energy calculation**

The folding energy was calculated through the equation:

$$\Delta G_0 = -k_BT\ln\left(\frac{p_{fold}}{p_{unfold}}\right) + \int_0^F (x_0(f) - x_u(f))df$$

The error of folding energy calculation came from two parts: one is the standard error of probability ratio in the first term, recorded as $\delta_1$. Another is the force calibration error in magnetic tweezer experiments, which is ~10%, exists in the second term, recorded as $\delta_2$.

To estimate the value of $\delta_1$, the dwell times of unfold state and refold state were collected and kept in two sets $U_{fold}$ and $U_{unfold}$, each contains 132 elements. The average unfolding state dwell time was 7.56s, while the average folding state dwell time was 7.86s. A python code-based bootstrap error estimation was performed. In each simulation, 132 elements were selected with replacement from $U_{fold}$ and $U_{unfold}$, respectively. $G_0 = -k_B T l n\left(\frac{p_{fold}}{p_{unfold}}\right)$ was calculated accordingly, and average value were obtained from the 132 $G_0$values. 1000 simulations have been performed and the $\delta_1 = Var(G_0)$was calculated to be $0.26 k_B T$.

In force calibration, the force has a variance of ~10%. By error propagation calculation, $\delta_2$ was calculated to be $1.9\, k_b T$ . Because $\delta_2$ and $\delta_1$ are independent, the total error equals to $\sqrt{{\delta_2}^2 + {\delta_1}^2}$=1.91 $k_b T$


1 Wang, Y., Yan, J. & Goult, B. T. Force-Dependent Binding Constants. *Biochemistry* **58**, 4696-4709, doi:10.1021/acs.biochem.9b00453 (2019).

2 Marko, J. F. & Siggia, E. D. Stretching dna. *Macromolecules* **28**, 8759-8770 (1995).

3 Winardhi, R. S., Tang, Q., Chen, J., Yao, M. & Yan, J. Probing Small Molecule Binding to Unfolded Polyprotein Based on its Elasticity and Refolding. *Biophys J* **111**, 2349-2357, doi:10.1016/j.bpj.2016.10.031 (2016).

4 Bouchiat, C. *et al.* Estimating the persistence length of a worm-like chain molecule from force-extension measurements. *Biophys J* **76**, 409-413 (1999).

5 Hugel, T. *et al.* Elasticity of single polyelectrolyte chains and their desorption from solid supports studied by AFM based single molecule force spectroscopy. *Macromolecules* **34**, 1039-1047 (2001).
